# Genomic regions and candidate genes affect root anatomical traits in diverse rice accessions

**DOI:** 10.1101/2023.04.24.538142

**Authors:** Jenna E. Fonta, Phanchita Vejchasarn, Meredith T. Hanlon, Susan R. McCouch, Kathleen M. Brown

**Author notes:** Corresponding author: Kathleen M. Brown.

## Abstract

Root anatomical traits show significant variation among rice, *Oryza sativa* L., genotypes and are of interest for improving adaptation to a variety of edaphic, hydrological and nutritional environments in which rice is grown. However, they are difficult to measure and the genetic controls of these traits are not well understood in rice. We conducted genome-wide association (GWA) analyses using moderate- and high-density SNP panels on a diverse rice population to identify genomic regions and candidate genes that control root anatomical traits. We identified 28 genomic regions for metaxylem vessel area and number, root cross-sectional area, stele area, and aerenchyma area. One genomic region associated with metaxylem vessel number and two regions associated with three root thickness-related traits, stele area, root cross-sectional area and metaxylem vessel area, were supported by chromosome-specific GWA using a high-density SNP panel and are regarded as highly significant regions controlling trait variation. Candidate genes in these regions were related to cell differentiation, elongation and division, and secondary cell wall formation. For genomic regions identified in the *indica* subpopulation, haplotype variation and root anatomical phenotypes were associated with geographic distributions of the accessions, notably the presence of alternate alleles conferring larger diameter roots, stele, and metaxylem vessels in accessions from the *indica 2* and *indica 3* subgroups originating largely in south and southeast Asia. The identification of genomic regions and candidate genes related to root anatomical traits in a diverse panel of rice varieties deepens our understanding of trait variation and genetic architecture and facilitates the incorporation of favorable alleles into breeding populations.

Key message

Genomic regions and candidate genes associated with root anatomical traits were identified in *Oryza sativa* using genome-wide association analyses.

## Introduction

Rice is consumed daily by over half of the world’s population, and reduced rice production severely impacts food security of the most vulnerable communities (McLean *et al*., 2013). Abiotic stresses related to water and nutrient uptake are major constraints to rice production, particularly in low-input systems. Drought stress in rainfed systems currently reduces rice yields by up to 25% (Zhang *et al*., 2018), and crop yields are expected to become at least 30% more variable due to variations in precipitation and temperature around the globe (Ray et al., 2015). Interest in root-related trait variation is also of great importance in breeding rice varieties that perform well under the “Alternate Wetting and Drying” (AWD) management regime designed to reduce irrigation water consumption and lower methane emmissions in rice paddies.

Roots play an important role in drought stress resistance because they are the organ responsible for intercepting, absorbing, and transporting water and nutrients to the rest of the plant. Rice root architectural traits, such as root growth angle, significantly impact drought tolerance. QTLs and genes controlling root growth angle, e.g. *DRO1*, have been mapped and incorporated into rice breeding lines to confer drought tolerance (Uga *et al*., 2013; Kitomi *et al*., 2015). Deep-rooted genotypes have been selected for in IRRI’s drought breeding program for many years, which has contributed in part in to the success of the drought-tolerant varieties they have produced (Henry, 2013). Variation in root anatomical traits may also have a significant impact on drought and low nutrient stress tolerance (Lynch, 2013), but so far these traits have been less well characterized in rice, particularly in mature plants.

Root anatomical traits consist of measurable features of the root cortex, stele, and entire cross-section. Variation in these traits has a direct impact on root resource capture. Axial hydraulic conductance, the volume of water transported axially along a length of root over time, is affected by the size and number of metaxylem vessels per root (Tyree and Ewers, 1991), which is directly relevant to plant performance under water-limited conditions. Roots with smaller metaxylem vessel diameter may benefit performance under terminal drought by reducing hydraulic conductance to conserve soil water resources (Comas *et al*., 2013). Smaller xylem vessels can also reduce the risk of cavitation and collapse when soil water is less available (Sperry and Saliendra, 1994; Guet *et al*., 2015). In wheat, smaller metaxylem vessels in seminal roots improved yield under drought (Richards and Passioura, 1989), and in maize, genotypes with reduced metaxylem area and hydraulic conductance performed better under drought (Klein *et al*., 2020). Some reports show, however, that upland rice varieties growing in drier conditions have thicker roots with larger xylem vessels, which provide both penetrability into hard dry soils as well as greater water uptake capacity (Henry, 2013). The ideal phenotype for xylem traits likely depends on the agroecosystem and the interaction with other traits, including root architecture.

Decreasing the metabolic cost of root tissue per length of root via anatomical trait variation could also improve drought tolerance (Lynch *et al*., 2014). Reduced root cross-sectional area, increased aerenchyma area (large air-filled lacunae in the root cortex), and/or reduced stele area decrease the volume of respiring living tissue per length of root and contribute to reduced root metabolic cost. With a decreased cost per unit root length, roots should be able to use the conserved metabolic energy to grow longer, deeper nodal and lateral roots to reach water resources deeper in the soil. In maize and common bean, the relationship between root anatomical traits and root respiration rates, a measure of root metabolic cost, has been associated with plant performance (Jaramillo *et al*., 2013; Saengwilai *et al*., 2014; Castañeda *et al*., 2018; Strock *et al*., 2018). Rice has substantial genetic diversity for root anatomical traits (Terashima *et al*., 1987; Champoux *et al*., 1995; Kondo *et al*., 2000; Uga *et al*., 2008; Uga *et al*., 2009), and some genetic loci that control these quantitative traits have been identified in biparental mapping populations (Uga *et al*., 2008), though more well-defined loci and specific alleles are needed to efficiently utilize these traits in breeding programs.

*Oryza sativa* is an diverse species composed of five major subpopulations: *indica* (*IND*), *tropical japonica* (*TRJ*), *temperate japonica* (*TEJ*), *aus* (*AUS*), and *aromatic* (*ARO*). In addition, *admixtures* between groups (ADMX) are observed, and many of these subpopulations consist of geographically defined subgroups. *Indica* for example has been classified into three (or sometimes four) subgroups, *ind1(A & B)*, *ind2*, and *ind3*, which have distinct histories and geographic origins (Wang et al., 2018b). To achieve greater adaptation to AWD management systems and provide improved abiotic stress tolerance in low input systems, this diversity must be explored and utilized. The Rice Diversity Panels 1 and 2 (RDP1, RDP2) and the 3,000 Rice Genomes (3KRG) are collections of rice accessions that together represent the global diversity of rice, and excellent tools have been developed to explore intraspecies diversity in these panels (Li et al., 2014b; McCouch et al., 2016). RDP1 and RDP2 were genotyped with a 700K SNP array (HDRA) that provided high-quality SNP genotypes (∼1 SNP every 540 bp) that could be used to identify associations with traits of interest (McCouch *et al*., 2016). The 3KRG accessions were re-sequenced, generating a set of 4.8M SNPs (The 3000 Rice Genomes Project, 2014), which increased marker density to one SNP every 89 bp. Using imputation, Wang *et al*. (2018a) leveraged information provided by the 4.8M SNPs from the 3KRG to increase SNP density on the RDP1 and RDP2 accessions and developed a Rice Reference Panel (RICE-RP) with 4.8M SNPs that linked the 3KRG, RDP1 and RDP2 for greater power in genomic exploration. In this study we investigated the genetic architecture of root anatomical traits in diverse rice genotypes from the RDP1 panel. We conducted genome-wide association (GWA) analysis using moderate marker density (HDRA and the RICE-RP pruned into subsets) to reduce error through multiple testing and high marker density (RICE-RP chromosome-specific) to enhance the resolution of GWA peaks to ultimately identify candidate genes controlling rice root anatomical traits.

## Materials and Methods

### Plant material and growing conditions

In this study, 335 rice (*Oryza sativa*) accessions from the Rice Diversity Panel 1 (RDP1) (Zhao et al., 2011) were grown in Penn State University greenhouses in University Park, PA (40°49’ N, 77°52’ W) and were phenotyped for root anatomical traits. Accessions represented all *O. sativa* subpopulations: *aus* (n=52), *indica* (n=67), *temperate japonica* (n=74), *tropical japonica* (n=80), *aromatic* (n=11), and admixed accessions (n=51, Supplemental Table 1). Three replications were grown per accession, and replications were staggered in time. Plants were grown in 10.5 L pots (21 cm x 40.6 cm, top diameter x height, Nursery Supplies Inc., Chambersburg, PA, USA) filled with growth medium consisting of 40% v/v medium size (0.3-0.5 mm) commercial grade sand (Quikrete Companies Inc., Harrisburg, PA, USA), 60% v/v horticultural vermiculite (Whittemore Companies Inc, Lawrence, MA, USA), and solid-phase buffered phosphorus (Al-P, prepared according to Lynch *et al*. 1990) providing a constant availability of high (100 µM) phosphorus concentration in the soil solution. Each pot was direct-seeded with three seeds, and after 7 days plants were thinned to one per pot. Plants were irrigated once per day via drip irrigation with Yoshida nutrient solution (Yoshida *et al*., 1976), which was adjusted to pH 5.5-6.0 daily.

### Root anatomy phenotyping

At the leaf 8 stage, root systems of each plant were excavated, washed with water, and preserved in 70% ethanol in water for later sampling. Two nodal roots approximately 20 cm long were sampled from each root system. Preserved root tissue at 15 cm from the root tip was freehand sectioned under a dissection microscope (SMZ-U, Nikon, Tokyo, Japan) at 4x magnification using Teflon-coated double-edged stainless-steel blades (Electron Microscopy Sciences, Hatfield, PA, USA). Transverse sections were stained with 0.05% Toluidine Blue. The three best root sections were selected from each nodal root sample, and images were captured using a Diaphot inverted light microscope (SMZ-U, Nikon, Tokyo, Japan) at 40x magnification equipped with digital camera (NIKON DS-Fi1, Tokyo, Japan). The two best images were chosen for quantitative analysis of root anatomical characteristics. The image analysis software Rice Root Analyzer (Taeparsartsit, 2013, unpublished) was used for analysis of root cross-section area (RXSA), aerenchyma area (AA), percentage of aerenchyma area per root cortical area (percAA), and number of metaxylem vessels (MXV). RootScan software (Burton *et al*., 2012) was used for analysis of root cortical area (TCA), stele area (TSA), and mean metaxylem vessel area (MXA) (Figure 1).

**Fig. 1.**
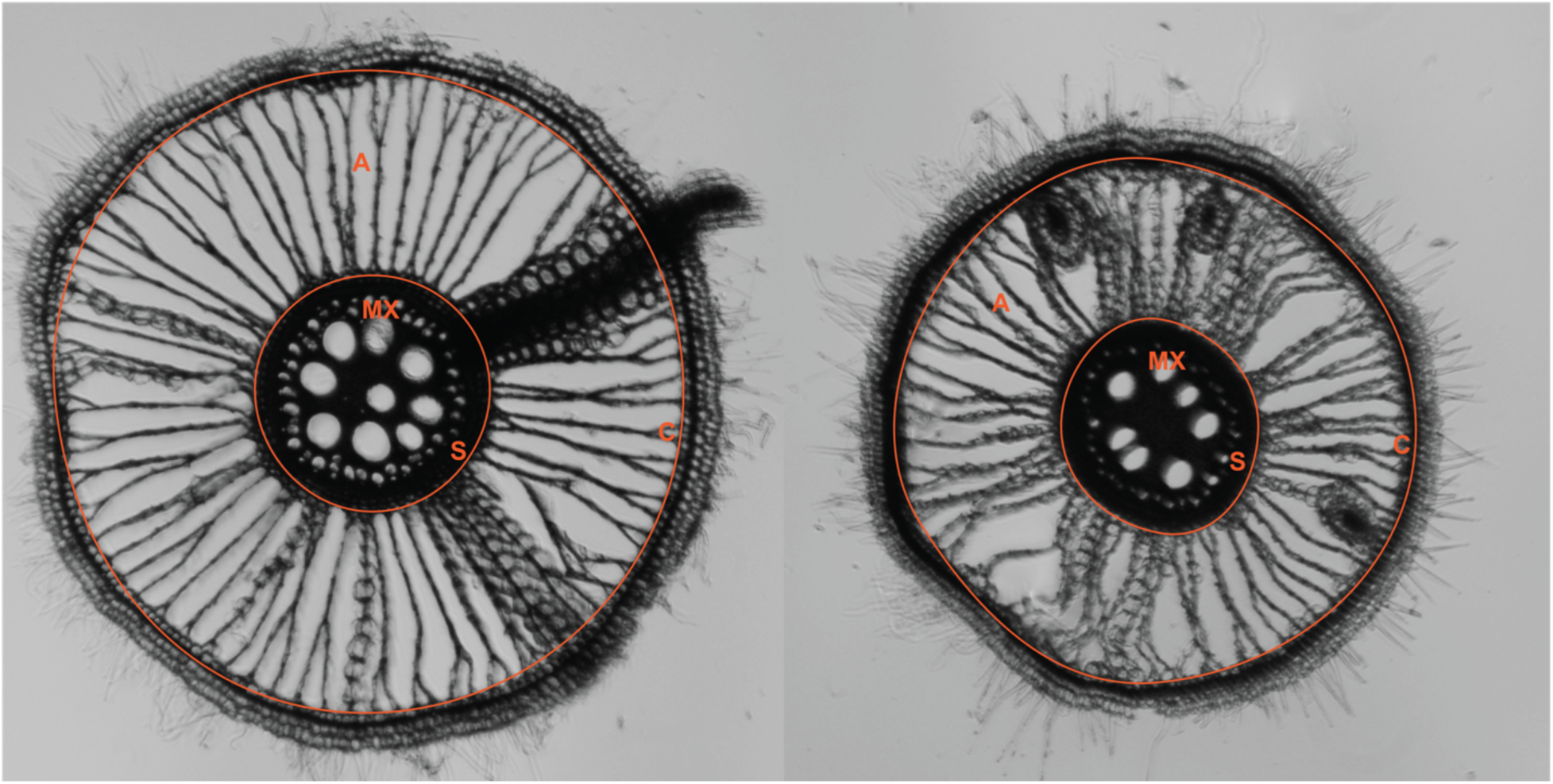
Rice root transverse cross-section images. Rice root transverse cross-sections showing variation in measured root anatomical traits. Representative metaxylem vessels (MX) and aerenchyma lacuna (A) are labeled. Area within small circle represents stele area (S), and area between small and large circle denotes root cortical area (C)

Using R version 3.6.3. (R Core Team, 2016), histogram, density distribution, and violin plots of each trait were generated in the *ggplot2* package (Wickham, 2016), and correlation coefficients and plots were generated with the *corrplot* package (Wei and Simko, 2017). Broad sense heritability or repeatability was calculated for each trait as:

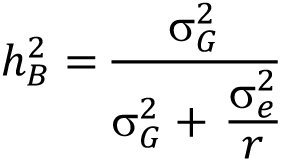

where α^2^_G_ represents genetic variance, α^2^_e_ represents residual variance, and r is the number of replicates (Fehr, 1991). Genetic coefficients of variation (GCV) for each root trait were calculated as:

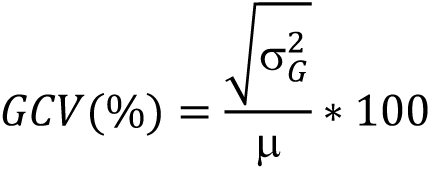

where μ is the population trait mean. Genetic and residual variances were determined for each trait by fitting linear mixed models using the restricted maximum likelihood (REML) method via R package *lme4* (Bates *et al*., 2015).

### Genome-wide association mapping with HDRA

Genome-wide association (GWA) analyses were conducted using the 700K SNP dataset (HDRA) (McCouch *et al*., 2016) and phenotypic data for five root anatomical traits (MXA, MXV, percAA, RXSA, and TSA) in all accessions (ALL) using three principle components to control for population structure, and within the four major subpopulations (*indica*, *IND*; *tropical japonica*, *TRJ*; *temperate japonica*, *TEJ; aus*, *AUS*). No principle components were used when performing subpopulation-specific analyses. Based on methods from McCouch *et al*. (2016), a linear mixed model in the gwas() function of the rrBLUP package (Endelman, 2011) was used for all association analyses. A minimum minor-allele frequency of 0.05 was used (min.MAF = 0.05) stipulating at least three minor allele count, and variance components were calculated once (P3D= TRUE). SNPs were filtered for heterozygosity within each subpopulation using plink (--hwe) at extreme p-vlaue thresholds to remove those with fewer than expected heterozygous SNPs. Significant SNPs were identified based on a *p*<0.0001 (-log_10_*p* > 4) and to control for Type I error, we classified genomic regions as significant only if at least three SNPs within 200 kb of each other were detected above the -log_10_*p* > 4 threshold. This was slightly more stringent than similar studies that have used the -log_10_*p* > 4 significance threshold (Zhao et al., 2011; Dimkpa et al., 2016; Kadam et al., 2017; Kadam et al., 2018). Manhattan plots were generated in the *ggplot2* package (Wickham, 2016).

### Genome-wide association mapping with RICE-RP

Using the imputed high density SNP dataset (4.8 million SNPs from the Rice Reference Panel (RICE-RP) (Wang *et al*., 2018a)), genome-wide associations were calculated for root anatomical traits in ALL, as well as within the major subpopulations (*IND*, *TRJ*, *TEJ*, *AUS)*. To avoid false-positives and reduce the computational load, SNPs were divided into 6 genome-wide subsets with non-orthogonal replacement based on step sizes of 5 and varying window sizes (25 or 50) and VIF/r^2^ thresholds (2/0.5 and 1.5/0.2) using the *indep* and *indep_pairwise* functions in plink1.9 (Chang *et al*., 2015) (Supplemental Table 2). Associations were calculated using gwas() in rrBLUP according to the same parameters as with HDRA (described above). Genomic regions were classified as in HDRA. As a further control for Type I error, we identified genomic regions that were significant in at least six of the seven HDRA and RICE-RP subset GWA analyses and required that the most-significant SNP (MS-SNP) was significant above a - log_10_*p* > 6 threshold. Genes within these regions were retrieved using Bioconductor tools in R (Huber *et al*., 2015).

Local linkage around genomic regions were assessed using the high-density RICE-RP dataset with the method described by Wang *et al*. (2017), though the critical r^2^ threshold was determined as the 75^th^ percentile of distribution of pairwise r^2^ across the chromosome instead of the 95^th^ percentile. Haplotypes were determined within select genomic regions using the most significant SNPs in the region with priority given to SNPs that were within gene models and that induced nonsynonymous mutations. Similar SNP genotypes were clustered using K-means clustering via the *pheatmap* v1.0.12 package (Kolde, 2019) and *factoextra* package v1.0.7 (Kassambara and Mundt, 2019) in R.

## Results

### Root anatomical variation in RDP1

Significant variation was detected for root anatomical traits in the RDP1 (Supplemental Figure 1). When subpopulations were compared, *tropical-japonica* (*TRJ*) accessions had significantly greater mean metaxylem vessel area, metaxylem vessel number, root cross-sectional area, cortical area and stele area than other subpopulations, while *indica* (*IND*) and *temperate-japonica* (*TEJ*) accessions had the smallest mean metaxylem vessel number, root cross-sectional area, and stele area (Supplemental Figure 1). Frequency distributions for root anatomical traits in ALL and in *IND* are shown in Figure 2. Within ALL accessions, trait broad-sense heritability averaged 0.86, ranging from 0.92 for stele area (TSA) to 0.81 for both mean metaxylem vessel area (MXA) and number of metaxylem vessels (MXV); similar heritabilities were seen within subpopulations (Supplemental Table 3). Genetic coefficients of variation for root anatomical traits showed similar trends, with TSA having the highest and MXV having the lowest coefficient of variation in both ALL and across subpopulations (Supplemental Table 3).

**Fig. 2.**
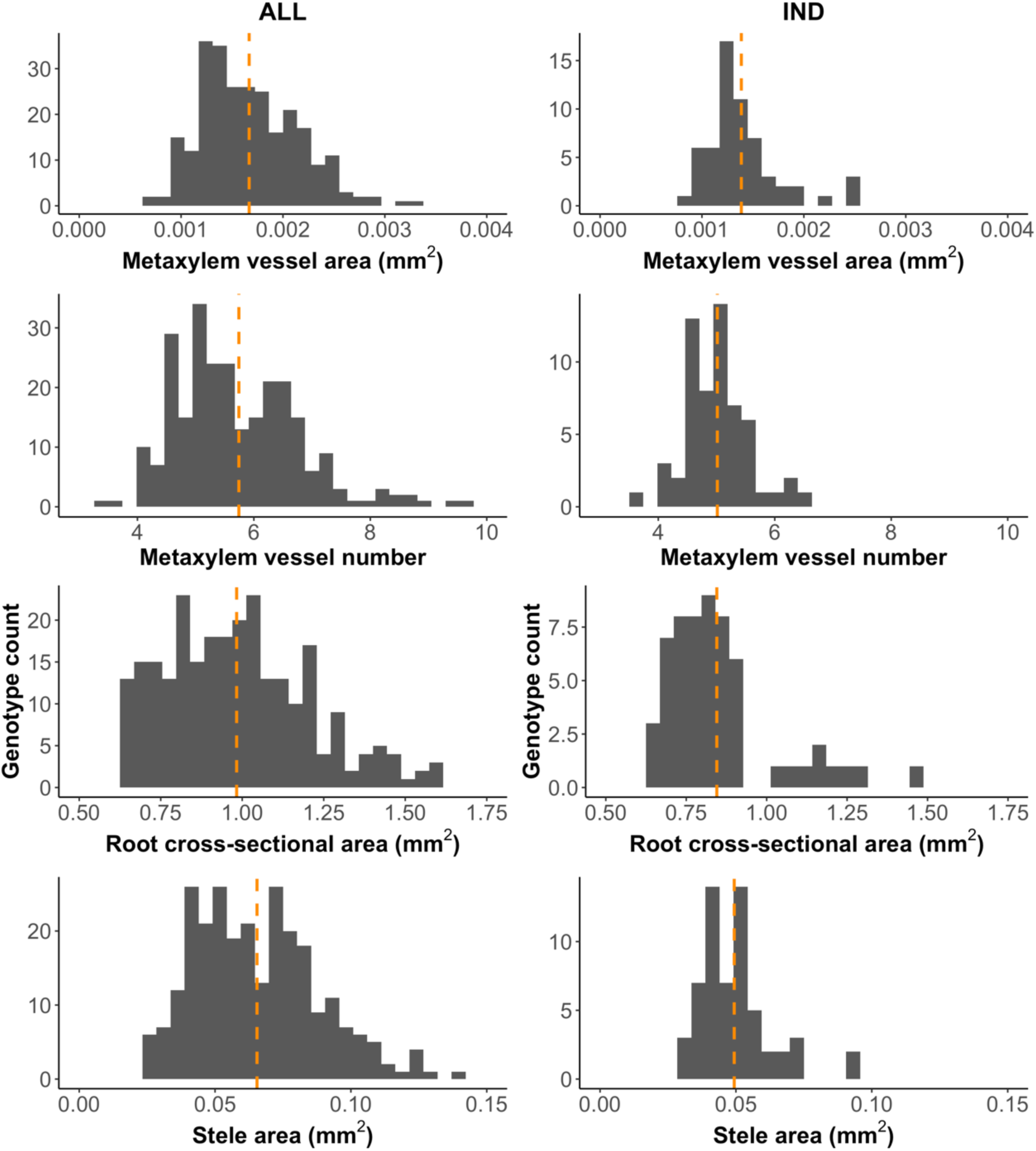
Distribution of root anatomical phenotypes in ALL and *IND*. Distribution of root anatomical traits in RDP1 accessions within combined subpopulations (n = 266) and in *indica* only (n = 59). Orange dotted lines represent median trait values among accessions

Root cross-sectional area (RXSA), cortical area (TCA) and stele area (TSA) were positively correlated in ALL and within each subpopulation (Figure 3, Supplemental Figure 2). The same three traits (RXSA, TCA and TSA) were positively correlated with MXA in *AUS, IND*, and *TEJ*, and with MXV in *TRJ* and *TEJ*. Negative correlations were observed between RXSA and percent aerenchyma area (percAA) in *AUS* and *TRJ*, indicating that thicker roots had proportionally less aerenchyma in those subpopulations., while there was a positive correlation in *TEJ* and no correlation in *IND*.

**Fig. 3.**
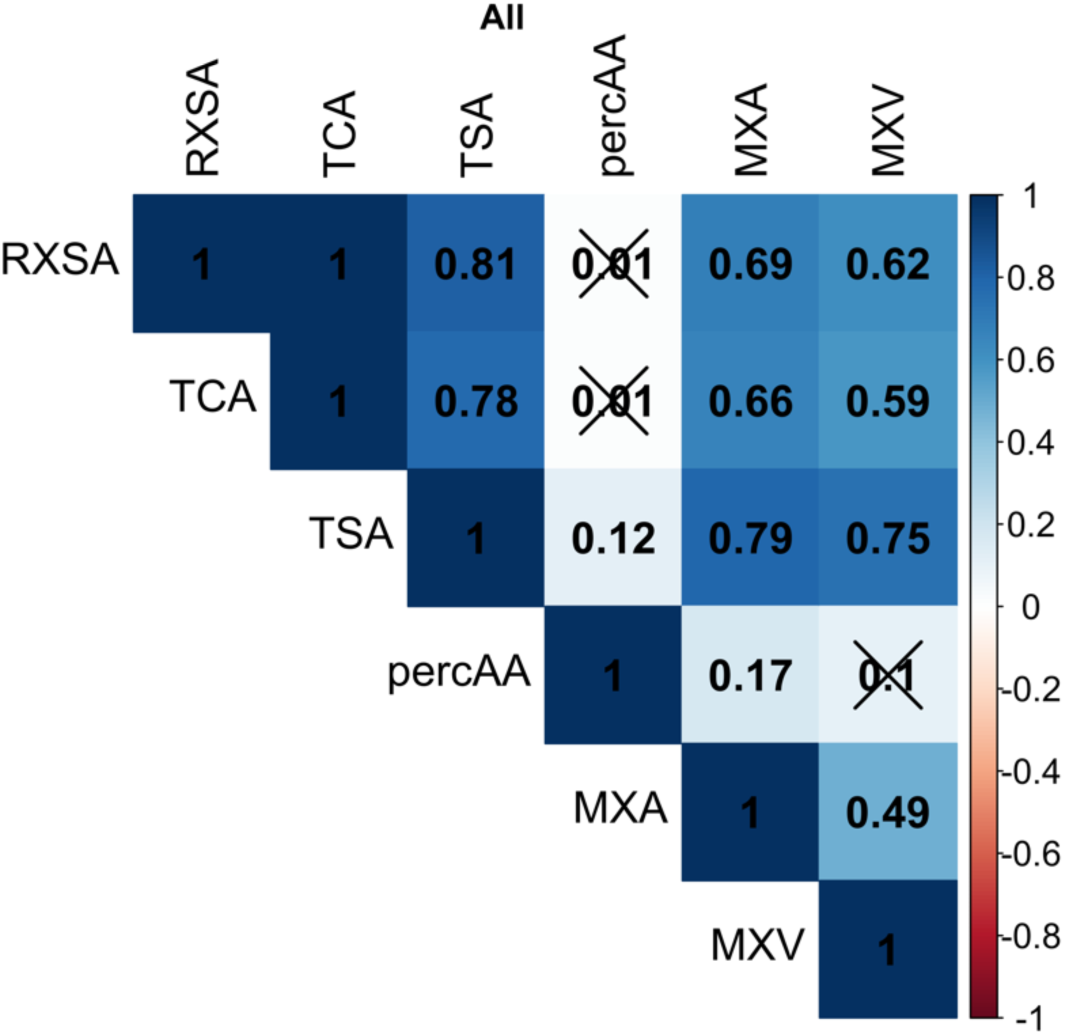
Correlation matrix of root anatomical traits within combined subpopulations (ALL). Each box shows the correlation coefficient, *r*, between each trait. “X”s indicate that the correlation was not significant at α = 0.1

### Genomic regions identified for root anatomical traits

A total of 28 genomic regions distributed on nine of the 12 chromosomes were significantly associated with the five root anatomical traits (Table 1). Six genomic regions (20%) were significant for multiple root anatomical traits, often for TSA and RXSA. Chromosome-specific association analysis using the high-density RICE-RP SNP set supported peaks identified in HDRA and the RICE-RP subsets in some, but not all cases (Figures 4-7).

**Fig. 4.**
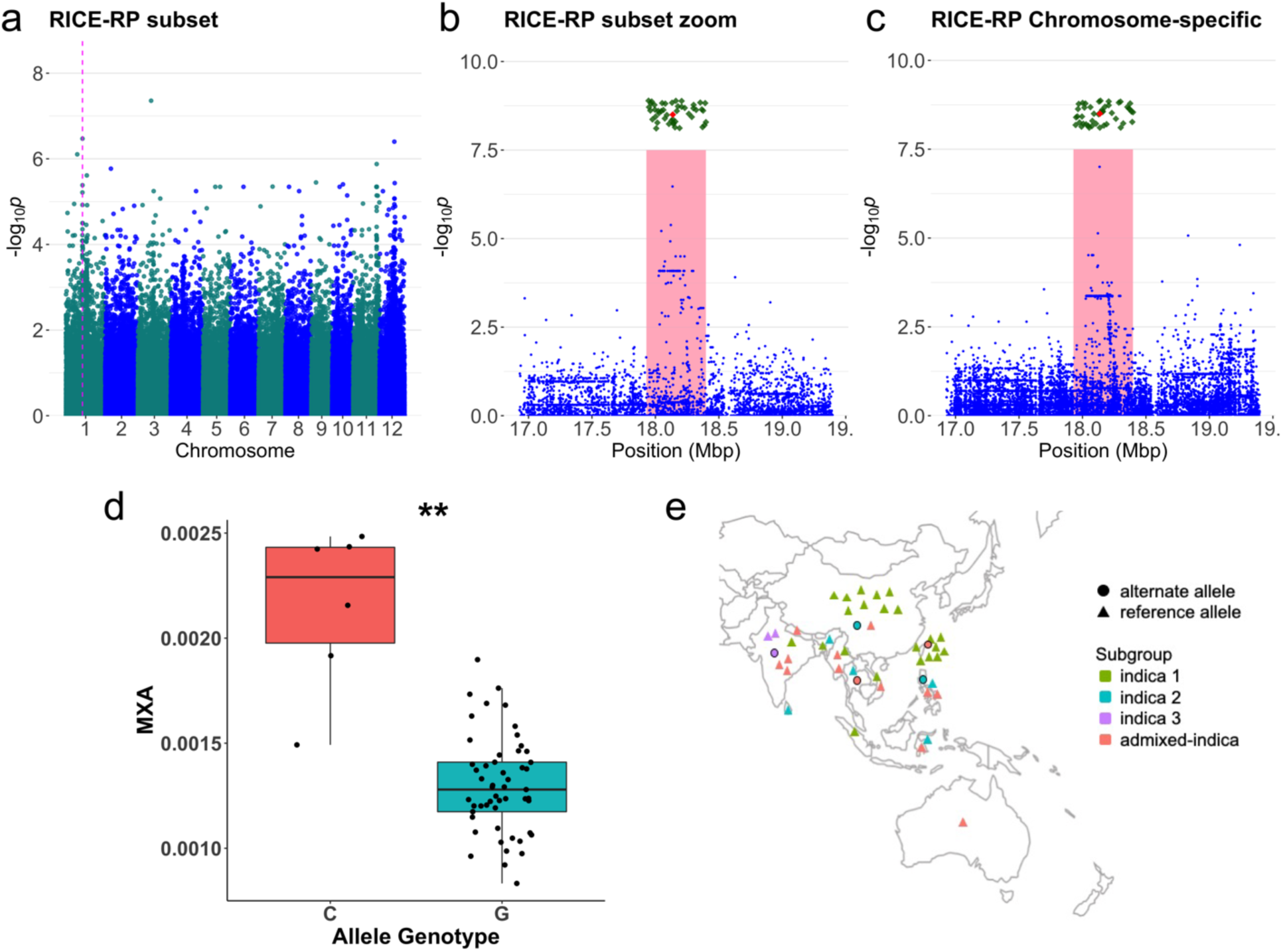
Manhattan plots, geographical distributions, and haplotype analysis of MXA *IND* Ch1 genomic region. Manhattan plots of metaxylem vessel area (MXA) GWA in the best RICE-RP subset a) across the whole genome, b) in zoomed view of the region, and c) in chromosome-specific GWA in *IND*. The whole genome plot shows the genomic region of interest at the pink dotted line. Zoomed views of chromosome 1 in the RICE-RP SNP subset and the RICE-RP chromosome-specific GWA show the genomic region in pink. Gene model positions are indicated as green diamonds, and red diamond indicates the gene model closest to the MS-SNP. d) Metaxylem vessel area (MXA) of RDP1 *IND* accessions with the major (G) and minor (C) allele of the MS-SNP. The significance level (*p-*value < 0.01**) is shown for the pair-wise t-test of MXA between alleles. e) Map showing country of origin of *IND* accessions with the allele that they carry at the MS-SNP of the genomic region

**Table 1.** Genomic regions for root anatomical traits in HDRA and RICE-RP subsets. Genomic regions are defined in the metaxylem vessel area (MXA), metaxylem vessel number (MXV), root cross-sectional area (RXSA), stele area (TSA), and percent aerenchyma area (percAA) GWA analyses of HDRA and RICE-RP subsets. Significance values (-log_10_*p*) and position of the most-significant SNP (MS SNP) in the genomic region are shown. Alternate allele effect represents a percent increase (+) or decrease (-) in trait average in accessions with the minor allele at the MS-SNP. Median significant SNP count in region among subsets indicates the median number of SNPs with -log_10_*p* > 4 within the significant GWA subsets. SNP subset count indicates the number of GWA subsets (min 0, max 7) where the genomic region met criteria to be significant. Regions in bold are further investigated in the text.

| Trait | Chr | Subpop | $-\log_{10}p$ of MS SNP | Mean alternate allele effect on phenotype | MS SNP position | Median significant SNP count in region among subsets | SNP subset count | Region start (bp) | Region end (bp) | Region size (kb) |
| --- | --- | --- | --- | --- | --- | --- | --- | --- | --- | --- |
| **MXA** | **1** | **IND** | **6.8** | **+ 23%** | **18134740** | **30** | **7** | **17929341** | **18397713** | **468** |
| **TSA** | **1** | **IND** | **5.7** | **+ 66%** | **18046581** | **28** | **7** | **17929341** | **18397713** | **468** |
| TSA | 1 | IND | 5.4 | + 43% | 22410649 | 5 | 6 | 22240519 | 22575982 | 335 |
| MXV | 2 | ALL | 5.0 | + 8% | 10385767 | 5 | 7 | 10119009 | 10711676 | 593 |
| MXV | 2 | ALL | 5.0 | + 16% | 27262183 | 4.5 | 6 | 27013624 | 27362183 | 349 |
| MXV | 3 | ALL | 5.1 | + 34% | 6562028 | 3 | 6 | 6327490 | 6662028 | 335 |
| MXV | 3 | ALL | 6.6 | + 34% | 7829729 | 7 | 6 | 7396992 | 7937705 | 541 |
| RXSA | 3 | IND | 6.0 | + 60% | 17257967 | 11 | 6 | 16997066 | 17529156 | 532 |
| RXSA | 4 | IND | 5.8 | + 55% | 5635275 | 3 | 6 | 5439264 | 5743290 | 304 |
| MXA | 6 | ALL | 5.9 | + 3% | 13390975 | 11 | 6 | 13168519 | 13577930 | 409 |
| percAA | 6 | ALL | 5.6 | - 15% | 26169601 | 5.5 | 6 | 26058095 | 26819617 | 762 |
| percAA | 8 | TEJ | 5.3 | + 13% | 14935615 | 5 | 6 | 14126366 | 15719032 | 1593 |
| RXSA | 10 | IND | 6.2 | + 49% | 11901414 | 3.5 | 6 | 11795359 | 12005976 | 211 |
| TSA | 10 | IND | 6.5 | + 76% | 11906904 | 4 | 6 | 11795592 | 12006904 | 211 |
| **MXV** | **10** | **ALL** | **6.3** | **+ 24%** | **15797419** | **11** | **7** | **15653828** | **16023182** | **369** |
| **MXA** | **11** | **IND** | **5.9** | **+ 69%** | **25839421** | **20** | **7** | **25485867** | **26131699** | **646** |
| **RXSA** | **11** | **IND** | **6.7** | **+ 57%** | **25886995** | **18** | **7** | **25500809** | **25986995** | **486** |
| **TSA** | **11** | **IND** | **7.1** | **+ 79%** | **25839421** | **21** | **7** | **25500809** | **25986995** | **486** |
| TSA | 11 | IND | 6.0 | + 80% | 27690597 | 19 | 7 | 27234698 | 27891367 | 657 |
| RXSA | 11 | IND | 6.0 | + 57% | 27609897 | 17 | 7 | 27509897 | 28133731 | 624 |
| **RXSA** | **12** | **IND** | **9.4** | **+ 59%** | **10232609** | **15.5** | **6** | **9888305** | **10760431** | **872** |
| **TSA** | **12** | **IND** | **7.9** | **+ 71%** | **10232609** | **17** | **6** | **9888305** | **10730198** | **842** |
| TSA | 12 | IND | 5.0 | + 97% | 12756769 | 5.5 | 6 | 12586741 | 13265300 | 679 |
| **TSA** | **12** | **IND** | **8.9** | **+ 50%** | **16290253** | **11** | **7** | **15515147** | **16800473** | **1285** |
| **RXSA** | **12** | **IND** | **9.5** | **+ 60%** | **16290292** | **15** | **7** | **15689547** | **17800299** | **2111** |
| **MXA** | **12** | **IND** | **6.4** | **+ 77%** | **16290253** | **17** | **7** | **16093794** | **17724267** | **1630** |
| **TSA** | **12** | **IND** | **8.1** | **+ 48%** | **17426742** | **12** | **6** | **17173010** | **17711872** | **539** |
| RXSA | 12 | IND | 7.4 | + 70% | 22709837 | 29 | 7 | 22148545 | 22955876 | 807 |

### Genomic region associated with MXA in IND on chromosome 1 demonstrates subgroup-specific variation

A 468-kb genomic region on chromosome 1 detected in the *IND* subpopulation was identified for MXA and TSA in all seven medium-density GWA analyses (Table 1). In this region, the most significant SNP (MS-SNP) was detected with -log_10_*p* > 6 in the genome-wide scans and showed a clear peak on the Manhattan plots (Figure 4a). Using the high-density RICE-RP SNP set to improve peak resolution, we performed chromosome-specific association analysis and the same MS-SNP was detected with greater significance (-log_10_*p* = 7) (Figure 4b-c). A total of 46 gene models are located in the region (green diamonds shown in Figure 4b-c, Supplemental Tables 4-5). Accessions carrying the alternate allele of the MS-SNP (n=6, C) have 65% greater MXA than those carrying the reference allele (n=53, G) (Figure 4d). Interestingly, lines carrying the alternate allele were classified as belonging to the *indica 2, indica 3* or *admixed indica* subgroups (Wang et al., 2017), while *indica 1* accessions all carried the reference allele at this locus (Figure 4e).

The MS-SNP was closest to a meristem structural organization gene associated with vesicle-mediated transport (Os01g0513700) (Supplemental Table 4). Other genes of interest within the genomic region are listed in Supplemental Table 4. Local linkage disequilibrium (LD) around this genomic region was predicted to extend 652 kb (17,925,833 bp to 18,577,790 bp), consistent with the genomic region (17,929,341 bp to 18,397,713 bp) estimated using the MXA GWA analysis (Supplemental Figure 3a).

### Genomic region associated with MXV in ALL on chromosome 10

A genomic region was identified on chromosome 10 for MXV in ALL accessions for all seven GWA analyses (Table 1). This region had an MS-SNP -log_10_*p* > 6 and a distinct peak on the Manhattan plots (Figure 5a, b), supported by chromosome-specific analysis with the high density SNP dataset (Figure 5c). A peak in the same region was detected in the *TRJ* subpopulation but the MS-SNP was located 203.5 kb downstream of the MS-SNP identified in ALL, and at lower significance (Figure 5d). Accessions carrying the alternate allele of the MS-SNP in ALL (n=22, T) had 24% greater MXV than those carrying the reference allele (n=280, C) (Figure 5g). Within the *TRJ* subpopulation, accessions carrying the alternate allele of the MS-SNP (n=3, T) had MXV values 29% greater than accessions carrying the reference allele (n=78, C) (Figure 5h).

**Fig. 5.**
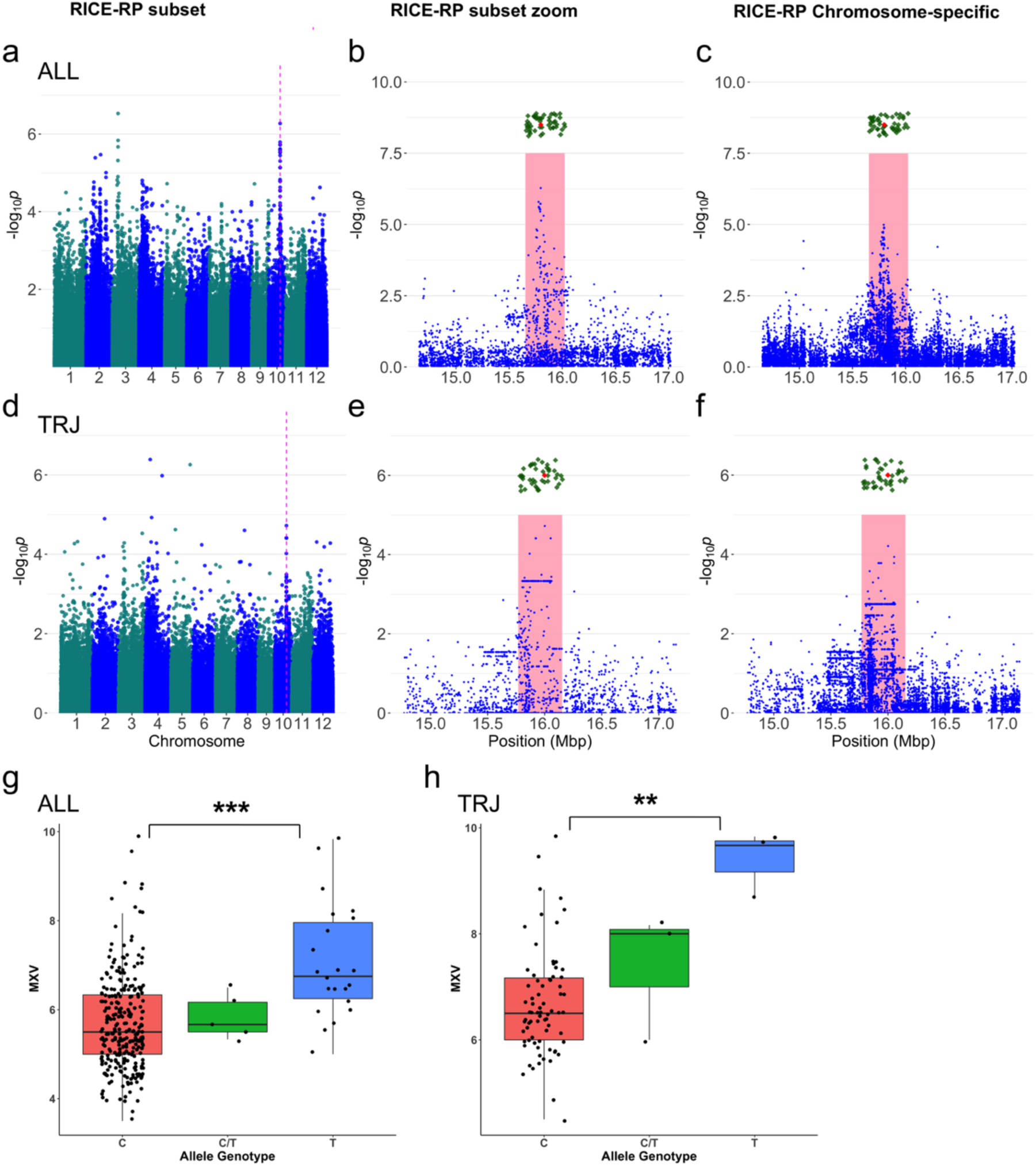
Manhattan plot and haplotype analysis of MXV ALL and *TRJ* Ch10 genomic region. Manhattan plots of metaxylem vessel number (MXV) GWA in the best subset a) across the whole genome, b) in zoomed view of the region, and c) in chromosome-specific GWA in ALL and in *TRJ* (d-f), respectively. The whole genome plot shows the genomic region of interest at the pink dotted line. Zoomed views of chromosome 10 in the best RICE-RP SNP subset and the RICE-RP chromosome-specific GWA show the genomic region in pink. Gene model positions are indicated as green diamonds, and red diamond indicates the gene model closest to the MS-SNP. Metaxylem vessel number (MXV) in ALL (g) and in *TRJ* (h) accessions with the major (C) and minor (T) allele of the MS-SNP. The significance level for ALL (*p-*value < 0.001***) and *TRJ* (*p-*value < 0.01**) are shown for the pair-wise t-test of MXV between alleles

Haplotypes were generated using significant SNPs from both MXV GWA analyses (Supplemental Figure 4a). Haplotypes found at high frequency in *TRJ* accessions (H5, H7, H9, H10, and H13) had significantly greater MXV than haplotypes found predominantly in *temperate japonica* (H1) or in *indica* and *aus* accessions (H3) (Supplemental Figure 4a-b). Accessions with the alternate allele of this MS-SNP did not overlap with accessions carrying the alternate alleles from the other genomic regions discussed here (Supplemental Table 6).

A total of 49 gene models are located in this region (Supplemental Table 4). The MS-SNP and five other SNPs from GWA analysis in ALL were located within a gene annotated for oxidoreductase activity (Os10g0439800) (Supplemental Table 4-5). Three other genes located within the genomic region were associated with cell wall development and eight were associated with transcription factor complexes or had DNA binding ability (Supplemental Table 4).

An extensive region of LD was detected around the MS-SNP (∼2 Mb, 14,563,701 bp to 16,587,483 bp) based on the critical r^2^ threshold at the 75^th^ percentile, which was much larger than the 369 kb genomic region determined from the MXV GWA analysis (15,653,828 to 16,023,182 bp) (Supplemental Figure 3b).

### Genomic region associated with TSA, RXSA, and MXA in IND on chromosome 11

A genomic region on chromosome 11 was identified in *IND* for TSA, RXSA, and MXA in all seven GWA analyses (Table 1). This region had an MS-SNP with -log_10_*p* > 7 for TSA (Figure 6a-b), and chromosome-specific analyses also supported this peak (Figure 6c). Peaks were also strong in RXSA and MXA (Figure 6d,e,g,h) and were supported by chromosome-specific analyses (Figure 6f,i). Accessions carrying the alternate allele of the TSA and MXA MS-SNP (n=4, A) had 79% greater TSA and 69% greater MXA than those carrying the reference allele (n=54, G) (Figure 6j,l). Accessions carrying the alternate allele of the RXSA MS-SNP (n=4, A) had 57% greater RXSA than those carrying the reference allele (n=55, C) (Figure 6j). Two accessions (Pappaku and Ming Hui) out of the four accessions carrying the alternate alleles for the TSA/ MXA MS-SNP also carried the alternate allele for the RXSA MS-SNP (45.6 kb away) (Supplemental Table 1). Though the number of accessions carrying the alternate allele in each case is low, it is interesting to note that accessions carrying the alternate alleles are classified as *ind2, ind3,* and ADMX*-IND*, while no accessions in the *ind1* subgroup carried the alternate allele at these MS-SNPs (Supplemental Table 6). A total of 38 genes are located in the genomic region (Supplemental Table 4-5).

**Fig. 6.**
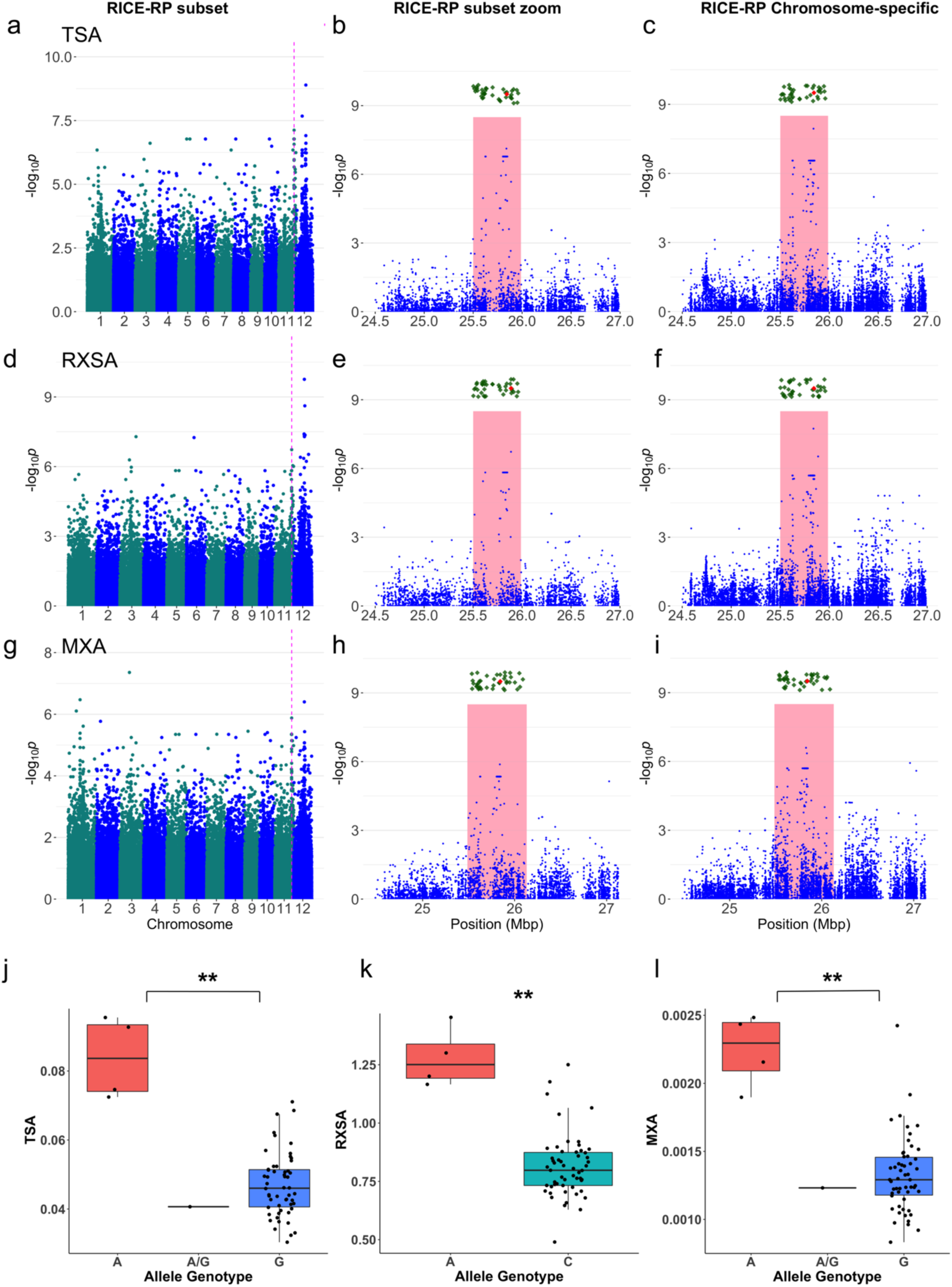
Manhattan plot and haplotype analysis of TSA, RXSA, and MXA *IND* Ch11 genomic region. Manhattan plot of stele area (TSA), root cross-sectional area (RXSA), and metaxylem vessel area (MXA) GWA in the best RICE-RP subset a,d,g) across the whole genome, b,e,h) in zoomed view of the region, and c,f,i) in chromosome-specific GWA in *IND*, respectively. The whole genome plot shows the genomic region of interest at the pink dotted line. Zoomed views of chromosome 11 in the best RICE-RP SNP subset and the RICE-RP chromosome-specific GWA show the genomic region in pink. Gene model positions are indicated as green diamonds, and red diamond indicates the gene model closest to the MS-SNP. j-l) Stele area (TSA), root cross-sectional area (RXSA), and metaxylem vessel area (MXA) of RDP1 *IND* accessions with the major and minor allele of the MS-SNPs. The significance level is shown for the pair-wise t-test of TSA (*p-*value < 0.01**), RXSA (*p-*value < 0.01**) and MXA (*p*-value < 0.01**) between alleles, respectively

Linkage around this genomic region was also assessed using a critical r^2^ threshold at the 75^th^ percentile of distribution of pairwise r^2^ across the chromosome. The linkage block covered 1.82 Mb (25,209,074 to 27,028,674 bp), and extended further on the distal side than the genomic region determined from the TSA GWA analysis (25,500,809 to 25,986,995 bp) (Supplemental Figure 3c).

### Two genomic regions associated with TSA, RXSA, and MXA in IND on chromosome 12

The first genomic region, situated at 9,888,305-10,730,198 bp on chromosome 12, was identified for TSA and RXSA in *IND* and was significant in six out of seven GWA analyses (Table 1). This region had an MS-SNP with - log_10_*p* > 7 for TSA and -log_10_*p* > 9 for RXSA and showed a clear peak on the Manhattan plots (Figure 7). Chromosome-specific GWA analysis also showed significant SNPs in this region (Figure 7c,f). Accessions carrying the alternate allele of the MS-SNP (n=4, A) had 71% greater TSA and 89% greater RXSA than those carrying the reference allele (n=54, G) (Figure 7j). It is interesting to note that two accessions, namely Ming Hui (*ind2*) and Pappaku (ADMX*-IND*) carried alternate alleles at this locus, and at both the RXSA MS-SNP and the TSA/MXA MS-SNP on chromosome 11 (Supplemental Table 1, 6). As in the other *indica* regions, all accessions carrying the alternate allele were in subgroups *ind2*, *ind3*, and ADMX*-IND* but not *ind1* (Supplemental Table 6).

**Fig. 7.**
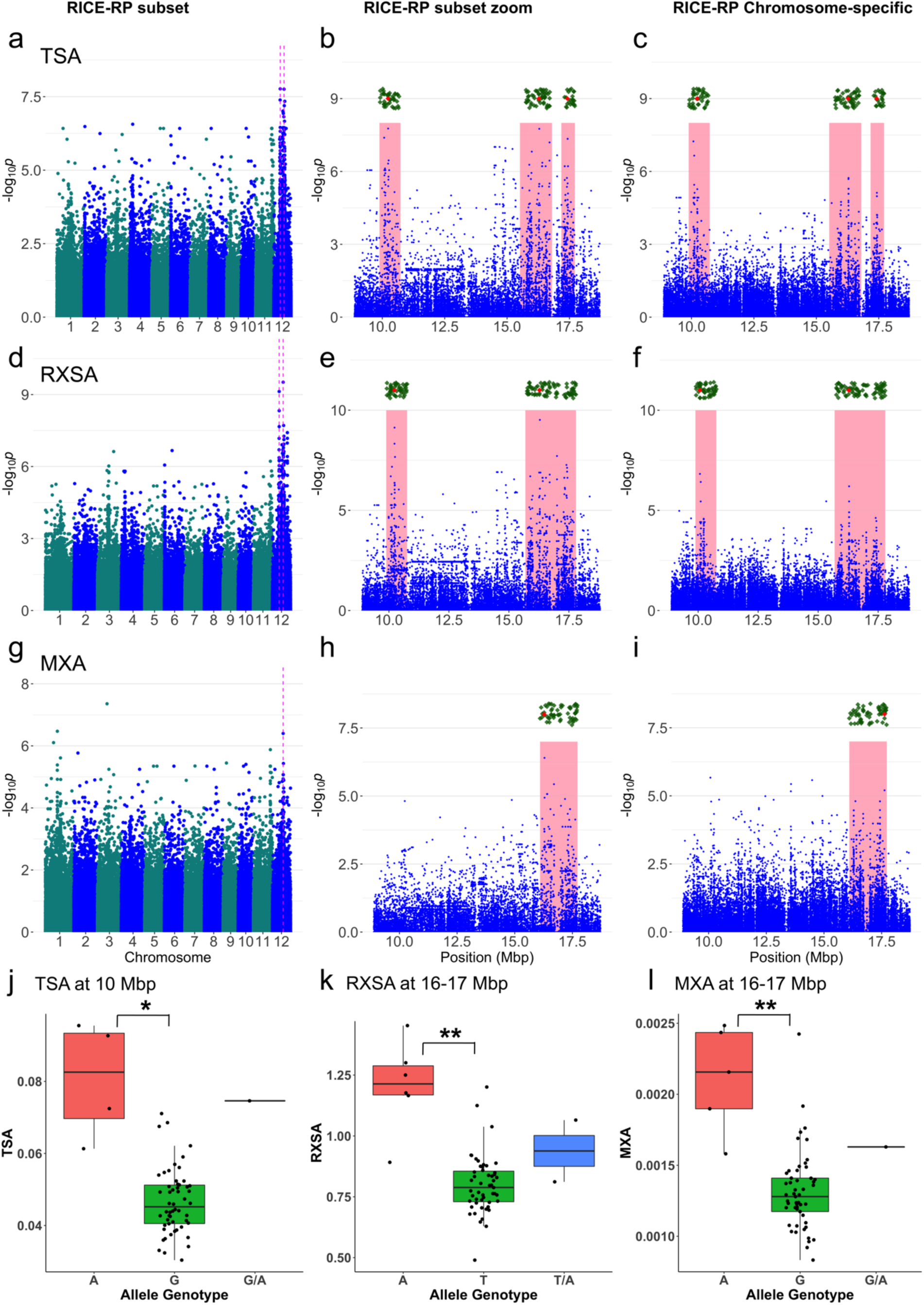
Manhattan plot and haplotype analysis of TSA, RXSA, and MXA *IND* Ch12 genomic region. Manhattan plot of stele area (TSA), root cross-sectional area (RXSA) and metaxylem vessel area (MXA) GWA in the best RICE-RP subset a,d,g) across the whole genome, b,e,h) in zoomed view of the region, and c,f,i) in chromosome-specific GWA in *IND*, respectively. The whole genome plot shows the genomic region of interest at the pink dotted line. Zoomed views of chromosome 12 in the best RICE-RP SNP subset and the RICE-RP chromosome-specific GWA show the genomic region in pink. Gene model positions are indicated as green diamonds, and red diamond indicates the gene model closest to the MS-SNP. j) Stele area (TSA) of RDP1 *IND* accessions with the major (G) and minor (A) allele of the MS-SNP at the 10 Mb peak. The significance level is shown for the pair-wise t-test of TSA (*p-*value < 0.05*) between major and minor allele accessions, respectively. k,l) Root cross-sectional area (RXSA) and metaxylem vessel area (MXA) of RDP1 *IND* accessions with the major and minor allele of the MS-SNP at their respective 16-17 Mb peaks. The significance level is shown for the pair-wise t-test of RXSA (*p-*value < 0.01**) and MXA (*p-*value < 0.01**) between major and minor allele accessions, respectively

There are 43 genes located in the region (Supplemental Table 4). The gene closest to the MS-SNP is a hypothetical gene coding region (Os12g0276550). Only one SNP fell within a gene model, a transcriptional repressor gene (Os12g0279100). Three other genes in the region were annotated for DNA-binding capacity and possible transcriptional regulation (Supplemental Table 4).

The linkage block extended 750 kb from 10,015,604 to 10,765,884 bp, slightly smaller than the genomic region determined from significant SNPs from the MXA GWA analysis, which was 842 kb wide from 9,888,305 to 10,730,198 bp (Supplemental Figure 3d).

The second genomic region on chromosome 12, located at 15,689,547-17,800,299, was identified for RXSA and MXA in *IND* for all seven GWA analyses (Table 1). In addition, TSA was significant for two overlapping regions at 15,515,147-16,800,473 and 17,173,010-17,711,872 (Table 1). This region had an MS-SNP -log_10_*p* > 9 for RXSA and showed a clear peak on the Manhattan plots (Figure 7), though chromosome-specific GWA analysis showed less support for these peaks (Figure 7c,f,i). Accessions carrying the alternate allele of the MXA MS-SNP and the first of the TSA MS-SNPs (n=5, A) had 77% greater MXA and 50% greater TSA than those carrying the reference allele (n=54, G) (Figure 7l). The accessions carrying the alternate allele at the RXSA MS-SNP (n=53, T) and the second TSA MS-SNP (n=6, A) averaged 60% and 48% greater RXSA and TSA, respectively, compared with accessions carrying the reference allele (Figure 7k, Table 1). As in the other *indica* regions, all accessions carrying the alternate allele were in subgroups *ind2*, *ind3*, and ADMX-*IND* but not *ind1*.

There are 78 genes located in the genomic region (Supplemental Table 4-5). The MS-SNP was closest to a putative phenylalanine amino-lyase protein (Os12g0461300), which is a putative transposon. Four significant SNPs fell with an oxidoreductase family gene (Os12g0464400). Within the genomic region, there were four genes annotated for cell fate determination or cell division, five for transcriptional regulation and DNA binding, and one for cellulose biosynthesis.

The linkage block in this genomic region extended 2.67 Mb, from 15,048,114 to 17,718,817 bp, and was larger than the genomic region defined in the RXSA and MXA GWA analyses (Supplemental Figure 3e).

### Accessions carry alternate alleles for large root anatomical traits at multiple genomic regions

A set of accessions in RDP1 carried the alternate alleles of the MS-SNPs for large TSA, RXSA, and MXA, while a separate set carried the alternate alleles for greater MXV (Supplemental Table 6). Accessions Ming Hui (*ind2*) and Pappaku (ADMX-*IND*) carried the alternate allele for all seven MS-SNPs for the TSA, RXSA, and MXA genomic regions on chromosome 1, 11, and 12. Accession JC149 (*ind3*) carried five alternate alleles for large TSA and MXA, and Bala (ADMX-*IND*) and Mudgo (*ind3*) carried four alternate alleles for large TSA, RXSA, and MXA (Supplemental Table 6). The TSA, RXSA, and MXA phenotypes for Ming Hui, Pappaku, JC149, Bala, and Mudgo were large compared to other *IND* accessions but were not the largest when compared to all accessions (Figure 2, Supplemental Figure 1, Supplemental Table 6). Accessions KU 115, Ligerito, and Pate Blanc MN 1 carried the alternate allele of the MS-SNPs for ALL and *TRJ* for greater MXV (Supplemental Table 6). These three accessions had very large values for MXV compared to other *TRJ* accessions and compared to all RDP1 accessions (Supplemental Figure 1, Supplemental Table 6).

## Discussion

### Root anatomical traits vary among subpopulations

Root anatomical traits, e.g. root cross-sectional area, cortical area, stele area, aerenchyma area, and metaxylem vessel area and number, vary significantly among rice accessions, highlighting the opportunities for both natural and artificial selection on root form and function, and underscoring the capacity of *Oryza sativa* to adapt to a variety of edaphic, hydrological and nutritional environments. In the RDP1, *TRJ* accessions had the thickest roots and consequently also had greater cortical area, stele area, metaxylem vessel area, and metaxylem vessel number compared to other subpopulations. This is supported by other studies that found *TRJ* accessions to have greater root cross-sectional areas than *IND* (Yadav et al., 1997; Gowda et al., 2011; Comas et al., 2013). As shown in these results as well as in other studies (Kadam *et al*., 2017; Hazman and Brown, 2018), root anatomical traits in rice are often correlated. Variation in root plasticity and root cross-sectional area has been shown to affect the ability of rice roots to penetrate hard soils (Gowda et al., 2011; Suralta et al., 2018) and root metabolic cost in other crops (Castañeda *et al*., 2018; Strock *et al*., 2018). Variation in metaxylem traits has been shown to affect growth, particularly under water-limited conditions in wheat (Richards and Passioura, 1989; Kadam et al., 2015) and maize (Klein *et al*., 2020). Identifying sources of root anatomical trait variation and markers and/or genes controlling those traits could greatly benefit breeding for abiotic stress tolerance in rice.

### Genomic regions control root anatomical traits

We conducted GWA analysis for rice root anatomical traits within and across subpopulations in the RDP1. Root anatomical traits had high broad-sense heritability, meaning trait variation was under genetic control, and this provided a strong foundation for the success of GWA analyses. Combined results from both HDRA and RICE-RP SNP sets identified 28 total genomic regions for metaxylem vessel area and number, stele area, root cross-sectional area, and aerenchyma area, six of which were significant for multiple root anatomical traits. Significant associations were detected in *IND*, *TEJ*, and ALL accessions in the RDP1. *IND* does not have greater metaxylem vessel area variation than other subpopulations, but greater SNP variation contributes to the ability to detect significant associations in the *indica* subpopulation. Due to the deep population structure in rice (McCouch *et al*., 2016; Gutaker *et al*., 2020), three principle components were used as co-variates in the GWA model when accessions from all subpopulations were analyzed together.

Using medium-density SNP sets from the HDRA and the RICE-RP strengthened our confidence in the genomic regions identified by GWA analysis. The HDRA captures approximately one SNP every 540 bp based on variation in a diversity panel consisting of 1500 accessions of *O. sativa* (McCouch *et al*., 2016) and the Rice Reference Panel (RICE-RP) provides one SNP approximately every 89 bp in the same panel, based on imputation from a set of 3,000 sequenced accessions from the IRRI gene bank (Li et al., 2014b; Wang et al., 2018b) where the imputation accuracy is >97% (Wang et al., 2018a). We utilized the RICE-RP by subsetting the 4.8 M SNPs into six non-orthogonal groups based on pruning by r^2^ and VIF thresholds. By independently pruning the number of SNPs in each RICE-RP subset, we removed SNPs in strong linkage with one another and obtained random collections of SNPs that were well distributed across the genome as the basis for GWA analysis. We also utilized the high marker density provided by the RICE-RP in chromosome-specific analyses to improve resolution of the peaks identified by GWA anayses using the HDRA and subsets of the RICE-RP data. Increasing marker density improved the resolution of significant peaks and genomic regions and provided insight into the extent of LD and the haplotype structure across these regions. This approach provided confidence that each of the regions reported in this study was significantly associated with the phenotype. As summarized in Table 1, we report genomic regions that were significant in at least six of the seven medium-density (HDRA and RICE-RP) GWA analyses, had highly significant MS-SNPs (-log_10_*p* > 6), showed up as strong peaks in Manhattan plots (median significant [-log_10_*p* > 4] SNP count > 10), had a demonstratable effect on phenotype, and/or were significantly associated with multiple anatomical traits.

### Candidate genes associated with larger metaxylem vessels and stele area on chromosome 1

We identified genes that mapped within each genomic region and listed those of particular interest in Supplemental Table 4, as well as provided an extensive list of all genes in each region in Supplemental Table 5. The MS-SNP in the genomic region on chromosome 1 associated with MXA and TSA in *IND* maps within the promoter region (3 kb upstream on reverse strand) of Os01g0513500, a hypothetical gene model. A second SNP, also significant for MXA, was found in the same gene model. This gene is expressed in root stele tissue according to online microarray expression data (RiceXPro). The Arabidopsis ortholog is expressed in root vascular tissue and root primordia (Maekawa et al., 2018; Choi et al., 2020). Further research is needed to determine whether these SNPs represent functional polymorphisms that alter expression of Os01g0513500 to increase MXA, or whether they are markers closely linked to other genes in the region that control MXA variation. The gene closest to the MS-SNP (forward strand) is Os01g0513700, annotated as “Similar to trafficking protein particle complex subunit 1, Sybindin-like protein family protein”. This gene has moderate expression in the division zone and elongation zone in root stele tissue according to RiceXPro expression data. GO terms related to this gene are vesicle-mediated transport, among others. The Arabidopsis ortholog, *BET5* (AT1G51160), is a TRAPP protein that is involved in vesicle trafficking relating to auxin distribution and cell wall formation in roots (Garcia et al., 2020). Three other genes in the genomic region of interest are related to cell-wall organization, and seven are related to transcriptional regulation. One gene that contains a significant SNP in the region and is cell wall-associated is Os01g0514700, a serine/threonine protein kinase-related protein. One or more of these genes may contribute to the variation in metaxylem vessel area associated with this genomic region.

### Candidate genes associated with metaxylem vessel number on chromosome 10

A prominent genomic region was associated with MXV on chromosome 10 in ALL accessions and in the *TRJ* subpopulation with a slightly shifted position of the MS-SNP. The MS-SNP in ALL and five other significant SNPs fall within a cytochrome P450 gene (Os10g0439800) annotated for oxidoreductase activity that is uncharacterized in rice or other species. Though this gene contains many significant SNPs that seem to be part of a group of *Japonica*-specific rare alleles on the dorsal side of the region, other genes in the region are better supported by the literature. One of the DNA-binding genes, Os10g0442600, annotated as “Similar to Cell division control protein 48-like protein e”, is orthologous to the *CDC48A* gene in Arabidopsis (AT3G09840), which encodes for a cell division cycle protein in which knock out mutants are defective in cytokinesis, cell expansion and cell differentiation (Park *et al*., 2008). This gene also contains SNPs significantly associated with MXV in the *TRJ* subpopulation. According to RiceXPro data, this gene is highly expressed in the division, elongation, and early maturation zones of rice root tips, particularly in the stele, meaning this gene could affect cell expansion and differentiation of immature metaxylem cells in these regions. One expansin gene (Os10g0439200) was also identified in this region and could have similar impacts on metaxylem cell size and number due to variations in cell expansion rates.

### Candidate genes associated with stele area, root cross-sectional area and metaxylem vessel area on chromosome 11

A strong genomic region on chromosome 11 was associated with TSA, RXSA, and MXA. Of the 38 genes in this region, nine contain significant SNPs (-log_10_*p* > 3). The gene closest to the MS-SNP (Os11g0648800) contains three significant SNPs, one of which is annotated as a nonsynonymous mutation, though the gene corresponds to a hypothetical gene model without characterization in rice or other species. Another gene in the region contains four significant SNPs and is annotated as an Na^+^/H^+^ antiporter, *OsNHX2*, (Os11g0648000), with moderate expression in root stele tissue and vascular bundles in the elongation and early maturation zone of rice based on RiceXPro and Fukuda *et al*. (2011). The Arabidopsis ortholog, *AtNHX2*, is a vacuolar-localized antiporter that is involved in growth regulation, stomatal closure, and reproduction (Bassil et al., 2011; Song et al., 2018). Double knock-out mutants *nhx1-nxh2* had significantly smaller cell size in Arabidopsis, particularly in hypotocyls and filaments, but also in root tissue (Bassil et al., 2011). It is therefore reasonable that alleles of this gene could have impacted the metaxylem cell size and cell size in other root tissues to alter RXSA and TSA.

### Two genomic regions found on chromosome 12 associate with stele area, root cross-sectional area and metaxylem vessel area

Two genomic regions located at 9-10 Mb and at 15-17 Mb on chromosome 12, respectively, are associated with TSA, RXSA, and MXA. One transcription factor in the region, a jumonji domain containing protein (JMJ702, Os12g0279100), contained one significant SNP and was annotated for histone modification and chromatin remodeling. A gene in the same family, *JM706*, has been shown to affect floral organ number and flower development via histone modification (Sun and Zhou, 2008; Chen et al., 2011). The Arabidopsis ortholog is known as *ETHYLENE INSENSITIVE 6 (EIN6), JUMONJI DOMAIN-CONTAINING PROTEIN 12 (JMJ12),* or *RELATIVE OF EARLY FLOWERING 6 (REF6)* (AT3G48430) and assists with histone mofications related to ethylene and brassinosteroid signaling, and to flowering time (Yu et al., 2008; Zander et al., 2019). Knock-out mutants *ref6* displayed reduced cell elongation as evidenced by short petioles in Arabidopsis (Yu et al., 2008). Alleles of *OsJMJ702* could also affect cell elongation in rice and alter TSA, RXSA, and MXA phenotypes.

The second genomic region on chromosome 12 at 15-17 Mb was not as well-defined in the chromosome-specific analysis, and the region was very large, including 78 gene models, making it difficult to identify likely candidate genes for controlling these root diameter-related traits.

This region overlaid closely with a previously identified large-effect rice root and drought-response QTL on chromosome 12, *qDTY_12.1_* (Bernier et al., 2009; Henry et al., 2014). Notably, NILs containing this QTL had greater transpiration efficiency and more conservative water usage than the parent line, with restricted water uptake early and greater uptake later during drought, along with more lateral root development (Bernier et al., 2009; Henry et al., 2014). In follow-up studies of genes controlling root branching density and drought response within a fine-mapped region of *qDTY_12.1_*, Dixit et al. (2015) identified a network of genes, including *OsARF1/24*, *OsCESA10, OsTDL1a*, *OsLGC1*, and *OsDEC* controlled by a master regulator *NO APICAL MERISTEM* (*OsNAM*), an NAC (NAM, ATAF1/2 and CUC2) family transcription factor. Root anatomical traits were not phenotyped in the Dixit *et al*. (2015) experiments, but since this region was highly significant in our results as well, these genes may have a major role in both root architectural and anatomical development in rice. NAC family proteins are known for regulating downstream genes associated with development (Xie et al., 2000; He et al., 2005), stress response (Fujita et al., 2004; He et al., 2005; Guo et al., 2017), secondary cell wall and lignin formation (Chai et al., 2015; Xu et al., 2015), and cell death (Wang et al., 2015), among others. Overexpression of *OsNAC5* led to increased root diameter and greater drought tolerance in Nipponbare (TEJ) rice (Redillas et al., 2012). Our genomic region included the same *OsNAM* gene (*OsNAC136*, Os12g0477400, LOC_Os12g29330) characterized within *qDTY_12.1_* by Dixit et al. (2015). This *OsNAM* may therefore control both root anatomical and architectural traits by regulating multiple pathways involved in these processes.

The effects of *qDTY12.1* and of our genomic region could result from action of other genes in the region in addition to or instead of *OsNAM*. *OsARF1/24* (Os12g0479400) are predicted transcriptional repressors, and *OsARF24* is known to act with *OsARF23* to affect cell growth and expansion in root tissue, as knock-out of both genes resulted in wavy growth of roots grown vertically (Li et al., 2014a). *OsARF1* knock-out mutants exhibited reduced root elongation (Song et al., 2009). These phenotypes suggest that this gene may be more involved in longitudinal cell expansion but could also impact radial variation in anatomical traits as measured in this study. *OsCESA10* (Os12g0477200) is thought to be involved in cellulose synthesis in cell walls, though since it has fewer functional domains than the other family members, its role may be weaker (Wang et al., 2010; Nakano et al., 2015). Alteration in the metaxylem cell walls by *CESA10* could still play a role in establishing the size of the vessels. *OsTDL1a* and *OsLGC1* were both annotated for cell fate determination, but their roles seem to be localized to reproductive tissues (Kusaba et al., 2003; Yang et al., 2016). *OsDEC* (Os12g0465700) plays a role in cytokinin signaling, phylotaxy, and reducing flowering delay under drought (Itoh et al., 2012; Sanchez et al., 2021), but its potential role in root anatomical development is not clear.

### Geographic distribution of accessions carrying alleles for larger metaxylem vessel, root cross-sectional, and stele areas in the indica subpopulation

In the RDP1, *indica* accessions can be classified into three subgroups which roughly correspond to geography. Of the *indicas* evaluated in this study, none of the accessions that were classified as *indica 1* (largely of Chinese and Taiwanese origin) carried the alternate allele at the MS-SNPs on chromosome 1, 11, or 12, which was observed primarily in accessions from South and Southeast Asia, identified with the *indica 2* and *indica 3* sub-groups. Historically rice has been widely grown in rainfed lowland and upland aerobic conditions in South and Southeast Asia and only relatively recently has it been widely cultivated in bunded flooded fields. On the other hand, flooded cultivation in anaerobic soils, i.e. the rice paddy system, was developed in China and has been common practice there for thousands of years (McLean *et al*., 2013). *Ind2* and *ind3* accessions carrying the alternate allele, with their larger metaxylem vessels, may be better adapted to aerobic soil conditions where less axial resistance to water movement provided by the large vessels could be advantageous, particularly when paired with deep roots for drought avoidance (Gowda et al., 2011). We expect that smaller vessels could be more adaptive in cultivars with shallow root systems grown in certain drought-prone environments because small vessels have a lower risk of cavitation and collapse and would therefore increase water use efficiency (Richards and Passioura, 1989; Sperry and Saliendra, 1994; Comas et al., 2013). The alternate allele accessions representing *ind2* and *ind3* also had greater root cross-sectional and stele area. Root diameter often decreases with drought stress, but the ratio of stele to root diameter increases, suggesting that prioritizing stele size could have advantages under drought, e.g. for root penetration (Henry et al., 2012; Hazman and Brown, 2018; De Bauw et al., 2019). Further research is needed to better understand the prevalence of the reference allele and the potential value of the alternate allele in the context of the other aspects of the root phenotype.

The genomic region on chromosome 1 was also significant for large stele area, which is highly correlated with metaxylem vessel area in *IND* and across all accessions. Similarly, correlations between these traits were found in a mapping population derived from an Indica X upland Japonica cross, and QTL mapping study of this population revealed one common QTL for both stele area and total metaxylem area on chromosome 9, along with another QTL for stele area and three other QTL for total metaxylem vessel area that were distinct (Uga et al., 2008). None of these QTL map within any of our genomic regions, but we found both distinct and overlapping genomic regions for area traits (MXA, TSA, RXSA) within the *indica* subpopulation. The persistence of the alternate alleles in *ind2* and *ind3* accessions could relate to adaptive advantages conferred by greater root, stele and/or metaxylem size in certain agroecologies of Southeast Asia.

### Leveraging SNP density to study genetic architecture of root anatomical traits

Using moderate- and high-density SNP sets to perform association analysis on accessions representing the diversity in *O. sativa*, we identified genomic regions and candidate genes that likely influence root metaxylem vessel area and number, root stele area, and root cross-sectional area. The strongest support is provided for three genomic regions (MXA *IND* Ch1; MXV ALL Ch10; TSA, RXSA and MXA Ch11); the other regions on chromosome 12 are strongly supported by GWA analyses, but less so by chromosome-specific analyses. This suggests that genes in these three regions explain only part of the phenotypic variation (23-79%) associated with the relevant traits and that other regions in the genome may interact with this region to control these anatomical traits.

Five accessions (Pappaku, Ming Hui, JC149, Bala, and Mudgo) carried the alternate alleles for at least four of the MS-SNPs associated with large stele, root cross-sectional, and/or metaxylem vessel area. Three accessions (KU 115, Ligerito, and Pate Blanc MN 1) carried the alternate alleles for the MS-SNPs for the ALL and *TRJ* genomic regions for large metaxylem vessel number. It is interesting to note that no accession carried all alleles for large root diameter (TSA, RXSA, MXA) and for large metaxylem vessel number. The anatomical traits related to root diameter are likely under separate genetic control than metaxylem vessel number, and the combination of large metaxylem vessel area and number may never be optimimal for physiological reasons related to root cost and efficient water conductance. These accessions may be good sources of alleles for increasing metaxylem vessel, stele, and root cross-sectional areas in rice breeding programs. Further genetic dissection of these loci might make use of these accessions that carry multiple alternate alleles as parents in crosses or as templates for gene validation via gene editing.

This study detected several alleles with large effects on root anatomical traits in rice. These alleles are globally rare but locally common, a pattern that is consistent with the inbreeding habit of the species. Beneficial alleles for root anatomical traits identified in *IND* accessions originating from Southeast Asia offer a valuable starting point for further genomic analyses, as well as phenotypic and physiological characterization and are of potential interest for improving root traits related to water and nutrient uptake in breeding programs. Our work is merely the beginning, but provides solid preliminary identification of alleles and genomic regions associated with traits that are difficult to measure and have long been ignored. Further work is needed to validate the effect of these alleles on root anatomical phenotypes in diverse genetic backgrounds and to rigorously evaluate their potential contribution to adaptive performance in diverse field environments. Elucidating the genetic architecture of root anatomical traits can lead to a deeper understanding of the hidden half of plants, and ultimately promises to enable more efficient and targeted improvement of rice varieties to enhance their resilience in our continually changing climate.

## Funding

This work was supported by The Royal Thai Government Scholarship Program, by the USDA National Institute of Food and Agriculture Federal Appropriations under project PEN04732, and by the National Science Foundation Plant Genome Research Program Grant #1026555. Any opinions, findings, and conclusions or recommendations expressed in this publication are those of the authors and do not necessarily reflect the views of the USDA National Institute of Food and Agriculture.

## Conflicts of interest/Competing interests

The authors have no conflicts of interest to declare that are relevant to the content of this article.

## Availability of data and material

Genotypic data from the High Density Rice Array (HDRA) is available as described in McCouch et al. (2016) and the imputed high density SNP dataset (4.8 million SNPs) from the Rice Reference Panel (RICE-RP) is available as described in Wang *et al*. (2018a). Phenotypic data from this study is provided in Supplemental Table 1. The p-values generated from a representative GWA analysis using the HDRA SNP data in combination with the seven root anatomical phenotypes measured in the the current study are available from the corresponding author upon request.

## Authors’ contributions

K.M.B. and P.V.conceived and designed the research with assistance from S.R.M.; P.V. performed the greenhouse experiments and initial data analysis; M.T.H. assisted with initial data analysis and developed analytical methods for subsequent analysis; J.E.F. adapted analytical methods for this data set; J.E.F. and K.M.B. analyzed the data; S.R.M. and M.T.H. contributed to data interpretation; J.E.F. and K.M.B. wrote the manuscript with input from S.R.M. and M.T.H.

## Supplemental Data Captions

**Supplemental Table 1.** Root anatomical phenotypes for all genotypes. Subpopulation (Subpop) is denoted as *indica* (*IND*), *temperate japonica* (*TEJ*), *tropical japonica* (*TRJ*), *aus* (*AUS*), *aromatic* (*AROMATIC*), and admixed genotypes (*ADMIX*). Rep indicates biological replications and Sub_rep indicates sampling replication with individuals. Root anatomical traits of root cross-sectional area (RXSA, mm^2^), cortical area (TCA, mm^2^), stele area (TSA, mm^2^), aerenchyma area (AA, mm^2^), percent aerenchyma area within the cortex (percAA, %), metaxylem vessel number (MXV), and mean metaxylem vessel area (MXA, mm^2^) are listed.

**Supplemental Table 2.** Comparison of SNP number in RICE-RP SNP subsets and HDRA used for root anatomy GWA analyses. SNPs within all accessions (ALL) and within *indica* (*IND*), *tropical japonica* (*TRJ*), *temperate japonica* (*TEJ*) and *aus* (*AUS*) are listed.

**Supplemental Table 3.** Root trait means, broad-sense heritability, and GCV by subpopulation. Means with 95% confidence interval, broad-sense heritability (h_B_^2^), and genetic coefficients of variation (%) of root anatomical traits among all accessions (ALL) and within each subpopulation. Root anatomical traits of root cross-sectional area (RXSA, mm^2^), cortical area (TCA, mm^2^), stele area (TSA, mm^2^), aerenchyma area (AA, mm^2^), percent aerenchyma area within the cortex (percAA, %), metaxylem vessel number (MXV), and mean metaxylem vessel area (MXA, mm^2^) are listed.

**Supplemental Table 4.** Genes of interest in selected genomic regions. Genes of interest in selected genomic regions. Nonsynonymous SNPs column indicates whether a gene contained a SNP that induced a nonsynonymous mutation in a coding region. Red font indicates gene closest to MS-SNP in genomic region.

**Supplemental Table 5.** All gene models in genomic regions. Red text indicates the gene closest to the MS-SNP. Bold text indicates that the gene contains a significant SNP (-log_10_*p* > 3) and purple fill indicates that the gene contains a significant SNP that induced a nonsynonymous mutation. Green fill in the GO name column indicates that the GO term is related to cell walls, blue fill indicates transcriptional or post-transcriptional regulation, and orange fill indicates cell differentiation or division.

**Supplemental Table 6.** RDP1 accessions with the Nipponbare (blue highlight) or alternate allele (orange highlight) of the MS-SNPs for the genomic regions of interest. Each of the first nine columns represents the genotype of the most significant (MS) SNP for genomic regions for metaxylem vessel area (MXA), stele area (TSA), and/or root cross-sectional area (RXSA) with their respective subpopulations. Each row represents genotypes at the MS-SNPs, accession information, and root anatomical phenotypes for each RDP1 accession. Only genotypes at MS-SNPs within the subpopulation where GWA was run are shown; otherwise ‘na’ is shown (eg. *IND* MS-SNPs do not have genotypes shown for non-*IND* accessions and *TRJ* MS-SNP does not have non-*TRJ* genotypes shown). Root phenotypes are colored based on the range of phenotypic values from smallest (blue) to largest (green). Rows are sorted within subpopulations by accessions with the greatest number of alternate alleles across the nine MS-SNPs.

**Supplemental Fig. 1** Distribution of root anatomical phenotypes. Distribution of root anatomical traits in RDP1 accessions within subpopulations (*AUS*, pink; *IND*, green; *TEJ*, blue; *TRJ*, purple). Diamonds represent trait median within subpopulation. Letters indicate significance groups at the α = 0.05 level determined by multiple comparisons tests using Tukey’s method

**Supplemental Fig. 2** Correlation matrices by subpopulation. Correlation matrices of root anatomical traits within *aus* (*AUS*), *indica* (*IND*), *tropical japonica* (*TRJ*), and *temperate japonica* (*TEJ*) subpopulations. Each box shows the correlation coefficient, *r*, between each trait. “X”s indicate that the correlation was not significant at α = 0.1

**Supplemental Fig. 3** Linkage plots of select genomic regions showing local linkage disequilibrium decay from the most significant SNP. Orange dotted line indicates the critical r^2^ threshold (75^th^ percentile); yellow curve shows local decay around the most-significant (MS)-SNP (red dot); green dotted line indicates linkage block bounds; pink dotted line indicates original genomic region bounds. a) MXA *IND* Ch1 MS-SNP is located at 18,134,740 bp, b) MXV ALL Ch10 MS-SNP is located at 15,797,419 bp, c) TSA *IND* Ch11 MS-SNP is located at 25,839,421 bp, d) TSA *IND* Ch12 at 10 Mb MS-SNP is located at 10,232,609 bp, e) TSA *IND* Ch12 at 16 Mb MS-SNP is located at 16,290,292 bp

**Supplemental Fig. 4** a) Haplotypes (H1-H13) in the metaxylem vessel number (MXV) genomic region on chromosome 10 within all accessions. Select significant SNPs in the HDRA and RICE-RP subset GWA are included. Nipponbare reference alleles are indicated in blue and alternate alleles are indicated in yellow. SNP physical positions (bp) and maximum significance values (-log_10_*p*) from MXV GWA in HDRA and RICE-RP subsets are listed. Red SNP position indicates the most-significant (MS) SNP in ALL, green SNP position indicates the MS-SNP from the *TRJ* GWA, bold SNP positions indicate that the SNP is within a gene model (listed below in the column) and purple bold SNP positions indicate SNPs that induce a nonsynonymous mutation with a gene model. The subpopulation identity of individual accessions with the haplotype are listed as well as the total number of accessions with the haplotype (n). Average metaxylem vessel number (MXV) is listed for each haplotype group. Letters indicate significance groups at the α = 0.05 level determined by multiple comparisons tests using Tukey’s method and *na* indicates that there is no test shown for that group because n = 1. b) MXV distribution of accessions within haplotypes with n > 3. Vertical lines indicate median values, and black points represent outliers beyond the 1.5x interquartile range

## Notes

### Competing Interest Statement

The authors have declared no competing interest.

