## Supplemental figures for "Genomic regions and candidate genes affect root anatomical traits in diverse rice accessions"

Fonta et al, Supplemental figures 1-6

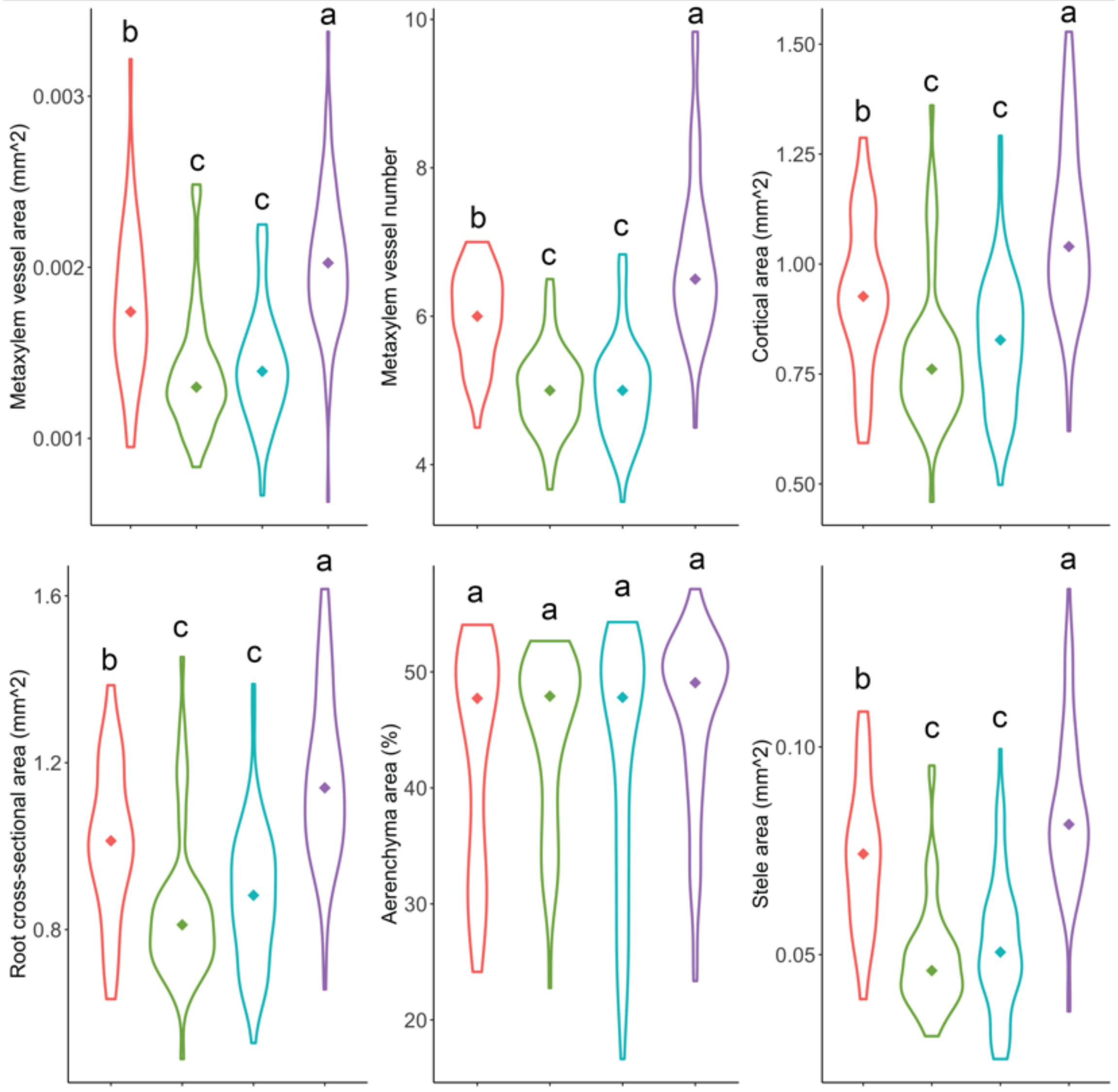


**Supplemental Figure 1** Distribution of root anatomical phenotypes. Distribution of root anatomical traits in RDP1 accessions within subpopulations (*AUS*, pink; *IND*, green; *TEJ*, blue; *TRJ*, purple). Diamonds represent trait median within subpopulation. Letters indicate significance groups at the $\alpha$ = 0.05 level determined by multiple comparisons tests using Tukey's method

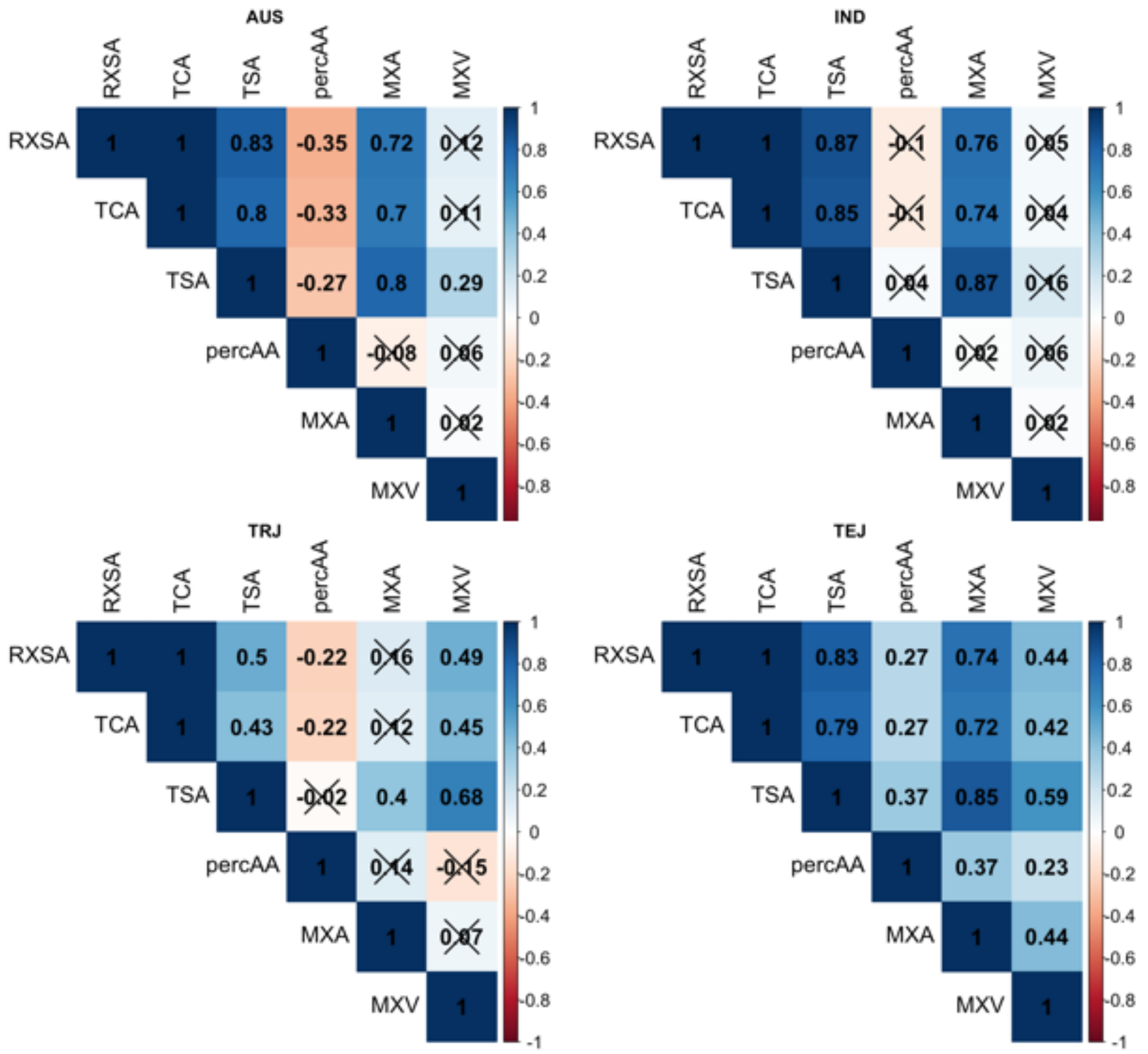


**Supplemental Fig. 2** Correlation matrices by subpopulation. Correlation matrices of root anatomical traits within *aus* (*AUS*), *indica* (*IND*), *tropical japonica* (*TRJ*), and *temperate japonica* (*TEJ*) subpopulations. Each box shows the correlation coefficient, *r*, between each trait. “X”s indicate that the correlation was not significant at $\alpha = 0.1$

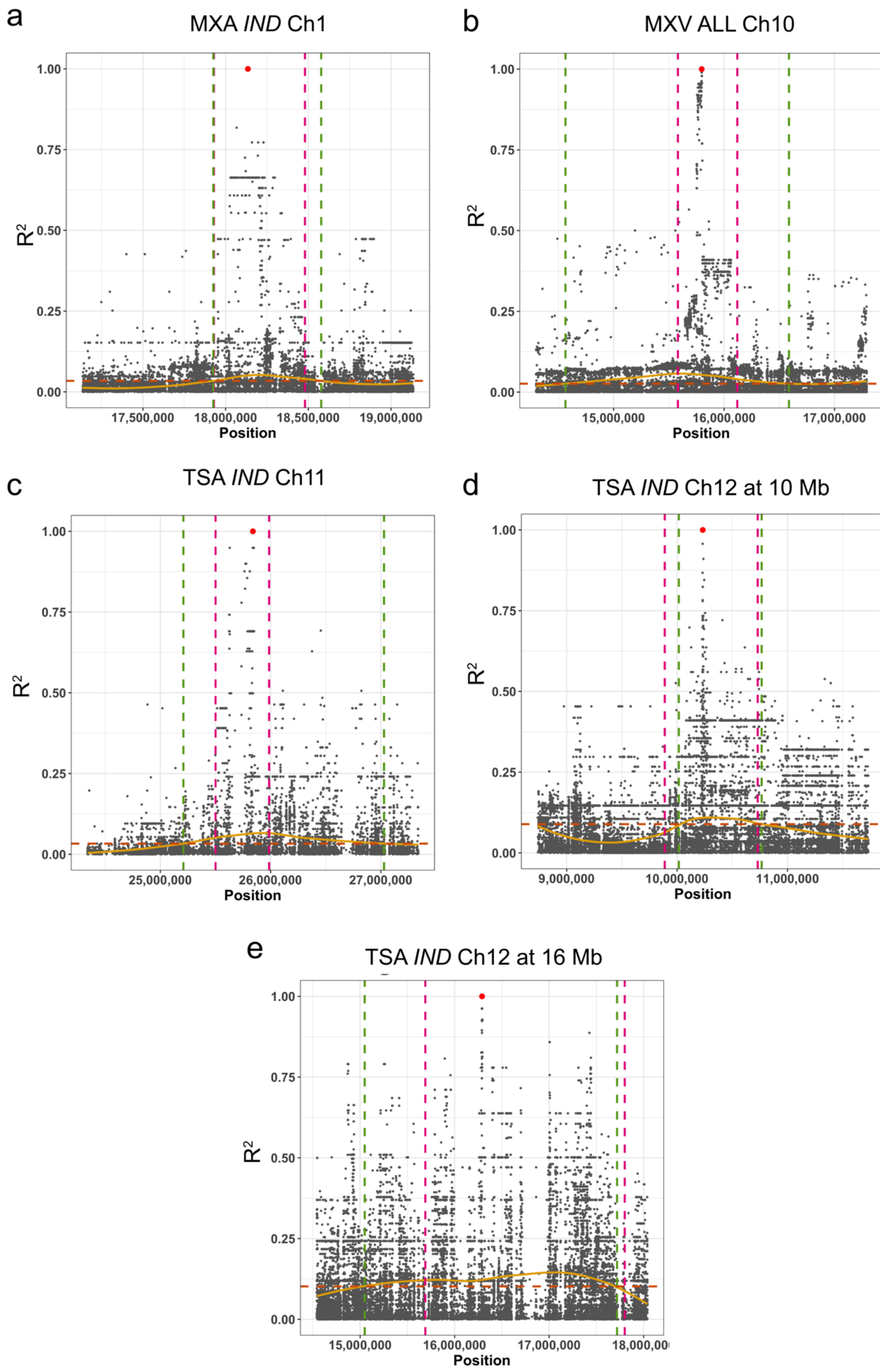


**Supplemental Fig. 3** Linkage plots of select genomic regions showing local linkage disequilibrium decay from the most significant SNP. Orange dotted line indicates the critical $r^2$ threshold (75th percentile); yellow curve shows local decay around the most-significant (MS)-SNP (red dot); green dotted line indicates linkage block bounds; pink dotted line indicates original genomic region bounds. a) MXA *IND* Ch1 MS-SNP is located at 18,134,740 bp, b) MXV ALL Ch10 MS-SNP is located at 15,797,419 bp, c) TSA *IND* Ch11 MS-SNP is located at 25,839,421 bp, d) TSA *IND* Ch12 at 10 Mb MS-SNP is located at 10,232,609 bp, e) TSA *IND* Ch12 at 16 Mb MS-SNP is located at 16,290,292 bp a

| | 15790283 | **15796312** | **15797419** | **15797513** | **15798037** | **15915504** | **15923182** | 16001086 | **16019688** | AUS | IND | TRJ | TEJ | ARO | ADMX | n | MXV |
|---|---|---|---|---|---|---|---|---|---|---|---|---|---|---|---|---|---|
| -logP | 5.5 | 5.5 | 6.3 | 6.0 | 4.2 | 4.6 | 4.2 | 4.7 | 4.0 | | | | | | | | |
| H1 | T | A | C | G | C | C | C | C | A | | 1 | 4 | **63** | | 5 | 73 | 5.1 **b** |
| H2 | C | A | C | G | C | C | C | C | A | | | | | | 2 | 2 | 5.3 **b** |
| H3 | C | A | C | G | C | T | C | C | A | 44 | **58** | 5 | | 10 | 8 | 125 | 5.5 **b** |
| H4 | C | A | C | G | C | T | T | C | A | **2** | | | | | | 2 | 6.0 **ab** |
| H5 | C | A | C | G | T | T | C | C | A | | | **58** | 4 | | 10 | 72 | 6.4 **a** |
| H6 | C | A | C | G | T | T | T | T | A | | | | 1 | | | 1 | 5.5 *na* |
| H7 | C | A | C | G | T | T | C | C | G | | | 1 | | | | 1 | 7.0 *na* |
| H8 | T | G | C | G | C | C | C | C | A | | | | **2** | | 1 | 3 | 5.7 **ab** |
| H9 | T | G | T | A | C | C | T | C | G | | | **3** | | | | 3 | 7.4 **a** |
| H10 | T | G | T | A | C | T | C | C | A | | | **6** | | | 2 | 8 | 7.0 **a** |
| H11 | T | G | T | A | C | C | C | C | A | | | | 1 | | **3** | 4 | 6.3 **ab** |
| H12 | T | G | C | A | C | C | T | T | G | | | | | | 1 | 1 | 5.5 *na* |
| H13 | T | G | T | A | C | C | T | T | G | | | 3 | 1 | | 3 | 7 | 7.5 **a** |
| Gene | | Os10g0439800 | Os10g0439800 | Os10g0439800 | Os10g0439800 | Os10g0442600 | Os10g0442800 | | Os10g0444400 | | | | | | | | |

b

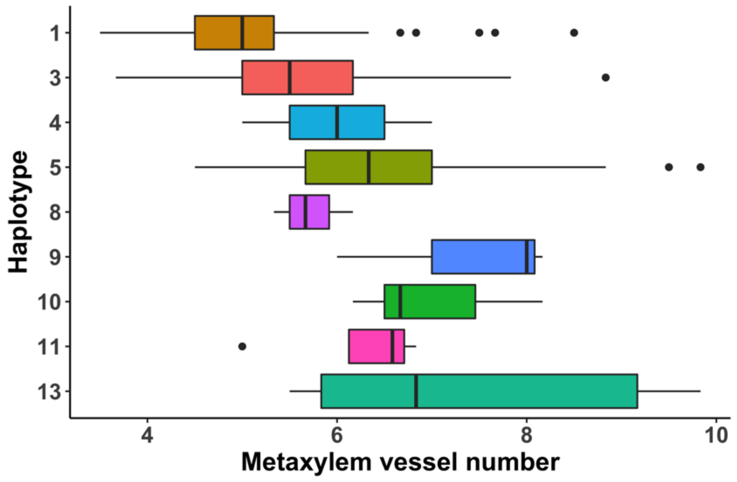


**Supplemental Fig. 4** a) Haplotypes (H1-H13) in the metaxylem vessel number (MXV) genomic region on chromosome 10 within all accessions. Select significant SNPs in the HDRA and RICE-RP subset GWA are included. Nipponbare reference alleles are indicated in blue and alternate alleles are indicated in yellow. SNP physical positions (bp) and maximum significance values (-$\log_{10}p$) from MXV GWA in HDRA and RICE-RP subsets are listed. Red SNP position indicates the most-significant (MS) SNP in ALL, green SNP position indicates the MS-SNP from the *TRJ* GWA, bold SNP positions indicate that the SNP is within a gene model (listed below in the column) and purple bold SNP positions indicate SNPs that induce a nonsynonymous mutation with a gene model. The subpopulation identity of individual accessions with the haplotype are listed as well as the total number of accessions with the haplotype (n). Average metaxylem vessel number (MXV) is listed for each haplotype group. Letters indicate significance groups at the $\alpha = 0.05$ level determined by multiple comparisons tests using Tukey's method and *na* indicates that there is no test shown for that group because n = 1. b) MXV distribution of accessions within haplotypes with n > 3. Vertical lines indicate median values, and black points represent outliers beyond the 1.5x interquartile range
